# Long acyl chain ceramides govern cholesterol and cytoskeleton dependence of membrane outer leaflet dynamics

**DOI:** 10.1101/663971

**Authors:** Anjali Gupta, Sneha Muralidharan, Federico Torta, Markus R. Wenk, Thorsten Wohland

## Abstract

The cellular plasma membrane composition and organization is crucial for the regulation of biological processes. Based on our earlier work showing that the same lipid probe, DiI, exhibits different dynamics in CHO-K1 and RBL-2H3 cells, we investigate the molecular factors that govern these differences. First, we determined that the cytoskeleton-interacting Immunoglobulin E receptor (FcεRI), which is abundant in RBL-2H3 but not in CHO-K1 cells, is not responsible for the DiI confinement found in RBL-2H3 cells. Second, lipid mass spectrometry of the plasma membrane of the two cells indicated differences in ceramide content, especially with long and very long acyl chains (C16 to C24). We, therefore, measure membrane dynamics by imaging total internal reflection fluorescence correlation spectroscopy in dependence on these ceramides. Our results show that C24 and C16 saturated ceramides uniquely alter the membrane dynamics by promoting the formation of cholesterol-independent domains and by elevating inter-leaflet coupling.

## INTRODUCTION

Among the factors contributing to the lateral heterogeneity in plasma membranes are lipid domains or “lipid rafts” and the cytoskeleton network in proximity to the plasma membrane inner leaflet^1–4^. To understand the complex membrane structure and dynamics, artificially reconstituted model membranes have provided important insights^5–15^. But they cannot recapitulate all physiologically relevant characteristics. For instance, in phase-separated model membranes, micron-sized domains of a specific phase (liquid disordered and liquid ordered) can be observed, while in cell membranes the size of the domains is below the diffraction limit^2^. In addition, about 30-50% of the area in a natural membrane is occupied by a range of structurally diverse proteins that influence the nanoscale organization of the membrane in critical ways^16–18^. Moreover, intact cell membranes exhibit phospholipid asymmetry which is challenging to mimic in model membranes, although recent progress in reconstituting asymmetric membranes has been made^19–21^. In asymmetric model membranes, the domain-forming lipids, such as sphingolipids, are localized in the outer leaflet while unsaturated lipids, such as PS, PI, or PA, which cannot form domains on their own, are localized in the inner leaflet of the membrane^22–24^. However, in cell membranes, a particular lipid may not be very strictly confined to one leaflet, and the proportion of lipids distributed in the two-leaflets can differ across cell types which can result in diverse membrane properties^25^. Recently, Li et al. have developed a method to estimate the proportion of lipids residing exclusively in the outer leaflet but this still needs to be utilized for the understanding of cell-type compositional differences^19^. Giant plasma membrane vesicles (GPMVs) preserve the compositional complexity of the cell membrane^26^ but they lack an actin cytoskeleton and the asymmetric organization of lipids and thus do not recapitulate the results of the original membranes. Therefore, for physiologically relevant results, it is necessary to analyze membrane dynamics in natural cell membranes.

In a recent study from our group, we compared the diffusive behaviour of two outer leaflet markers 1,1′-dioctadecyl-3,3,3′,3′-tetramethylindocarbocyanine (DiI-C_18_), an outer membrane free diffusion marker, and GFP-GPI, a marker of domain confined diffusion, across five cell types namely, CHO-K1, Hela, RBL-2H3, SH-SY5Y, WI-38^27^. We utilized imaging total internal reflection fluorescence correlation spectroscopy (ITIR-FCS) and the FCS diffusion law to obtain the diffusion coefficient (*D*) and diffusion law intercept (*τ_0_*), an indicator of membrane organization. Our results showed that DiI-C_18_ exhibits confined diffusion in RBL-2H3 cells at 298 K unlike in the other four cell lines tested where it shows free diffusion. Furthermore, we estimated the Arrhenius activation energy (E_*Arr*_) of diffusion, a determinant of molecular packing, for the markers across these cell types. The E_*Arr*_ of DiI-C_18_ in RBL-2H3 cell membranes was significantly higher compared to other cell lines and was comparable with the E_*Arr*_ of a cholesterol-dependent domain marker (GFP-GPI). Consequently, in RBL-2H3 cells DiI-C_18_ showed properties of domain confined diffusion and a weak dependence on the cytoskeleton. These observations imply a stronger transbilayer coupling in RBL-2H3 cells as compared to CHO-K1 cells, and indicate some DiI-C_18_/domain interactions in RBL-2H3 cell membranes. This study demonstrated that membrane lateral dynamics and organization varies across different cell types, and there is a differential strength of inter-leaflet coupling across the cell types^27^. We therefore endeavoured to identify which membrane components govern the lateral and transbilayer dynamics in cell membranes.

There is substantial evidence for the occurrence of transient nanodomains in the outer leaflet of cell membranes that can explain their lateral organization^14,28–39^. However, transbilayer coupling in cell membranes is not very well understood. One of the most important propositions to explain this phenomenon is the “picket fence model.” According to the “picket fence model,” membrane compartmentalization and transbilayer coupling can be mediated by transmembrane proteins, which are in contact with the actin cytoskeleton network^1^. In addition, reports are suggesting that long acyl chain or negatively charged lipids are responsible for transbilayer coupling. Experimental reports based on model membranes have shown that domains in the outer leaflet of the membrane can influence the organization of the membrane inner leaflet resulting in inner-leaflet domain formation^40,41^. Raghupathy et al. proposed that interaction of the actin cytoskeleton with the inner leaflet long acyl chain lipids is responsible for transbilayer coupling^42^. They observed that long acyl chain phosphatidylserine lipids in the inner leaflet interdigitate with the long acyl chain sphingomyelins in the outer leaflet of the membrane, thereby facilitating transbilayer coupling. It was also observed that the actin cytoskeleton forms domains in the membrane via their contacts with phosphoinositide lipids in a concentration-dependent manner^43^. Thus, both membrane lipids and transmembrane proteins can form ordered membrane domains and can mediate transbilayer coupling. Based on the observations mentioned earlier and existing literature^16,44–50,27^, RBL-2H3 cells are a potential model system to characterize the factors that cause transbilayer coupling.

In the current work, we study the effect of transmembrane protein density on DiI-C_18_ diffusion in RBL-2H3 cell membranes and how it influences the link of outer leaflet lipid dynamics with cholesterol and the cytoskeleton network. One of the most abundant transmembrane receptors in RBL-2H3 cells is the high affinity Immunoglobulin E receptor (FcεRI). FcεRI interact with the actin cytoskeleton and induce compartmentalization in the membrane^49,48^. Moreover, DiI-C_18_ co-distributes with the high density of FcεRI on RBL-2H3 cell membranes^44,45^. So, we reduced the density of FcεRI to determine if that weakens the connections of DiI-C_18_ diffusion with cholesterol and cytoskeleton. Furthermore, we compare the lipid composition of RBL-2H3 and CHO-K1 plasma membranes using mass spectrometry. Our results show that RBL-2H3 cell membranes possess significantly higher levels of ceramides and sphingomyelins. We therefore manipulate the levels of sphingolipids in the cell membrane and probe how the links of the outer leaflet lipid diffusion with cholesterol and the cytoskeleton are influenced. We show that the tuning of sphingolipid composition plays a crucial role in orchestrating the link of cholesterol and cytoskeleton with the outer leaflet lipid diffusion. Specifically, ceramides are sufficient to induce connections of the outer leaflet lipid diffusion with cholesterol and cytoskeleton in cell membranes. Furthermore, ceramides with saturated long and very long acyl chains show a greater tendency to form cholesterol independent domains and mediate cytoskeleton-induced domain formation in the outer leaflet.

## RESULTS

In this section we report on the dynamics and organization of cell membranes using two parameters measured by ITIR-FCS. The first is the diffusion coefficient (*D*) that reports on the mobility of molecules in the membrane and is inversely related to viscosity and which is expected to decrease with an increase of transient trapping of molecules in domains. The second parameter is the diffusion law intercept (*τ*_*0*_). The diffusion law, as explained in the Materials and Methods section, measures how the diffusion coefficient changes with the length scale, i.e. the size of the area, over which it is measured^51,52^. The diffusion law intercept is expected to be close to 0 for free diffusion. It is expected to be positive for transient entrapment (*τ*_*0*_ > 0.1s), e.g. in cholesterol dependent domains, and negative (*τ*_*0*_ < −0.1s) for hop diffusion, i.e. if molecular diffusion is hindered by a meshwork, e.g. the cytoskeleton (Figure 1*A*). In cases where a molecule is transiently trapped and at the same time is hindered by a meshwork, the intercept will be a weighted average and thus the absolute intercept value is not indicative of the diffusive mode. In these cases, we have recently shown that a decrease or increase in the intercept upon disruption of domain or the cytoskeleton is an indication for transient trapping or hop diffusion, respectively^53^. Finally, measurement errors for the diffusion coefficient on the same cell are typically 20% (Figure *S1D*) and we therefore will report changes in diffusion coefficient as significant only if they exceed this value.

**Figure 1:**
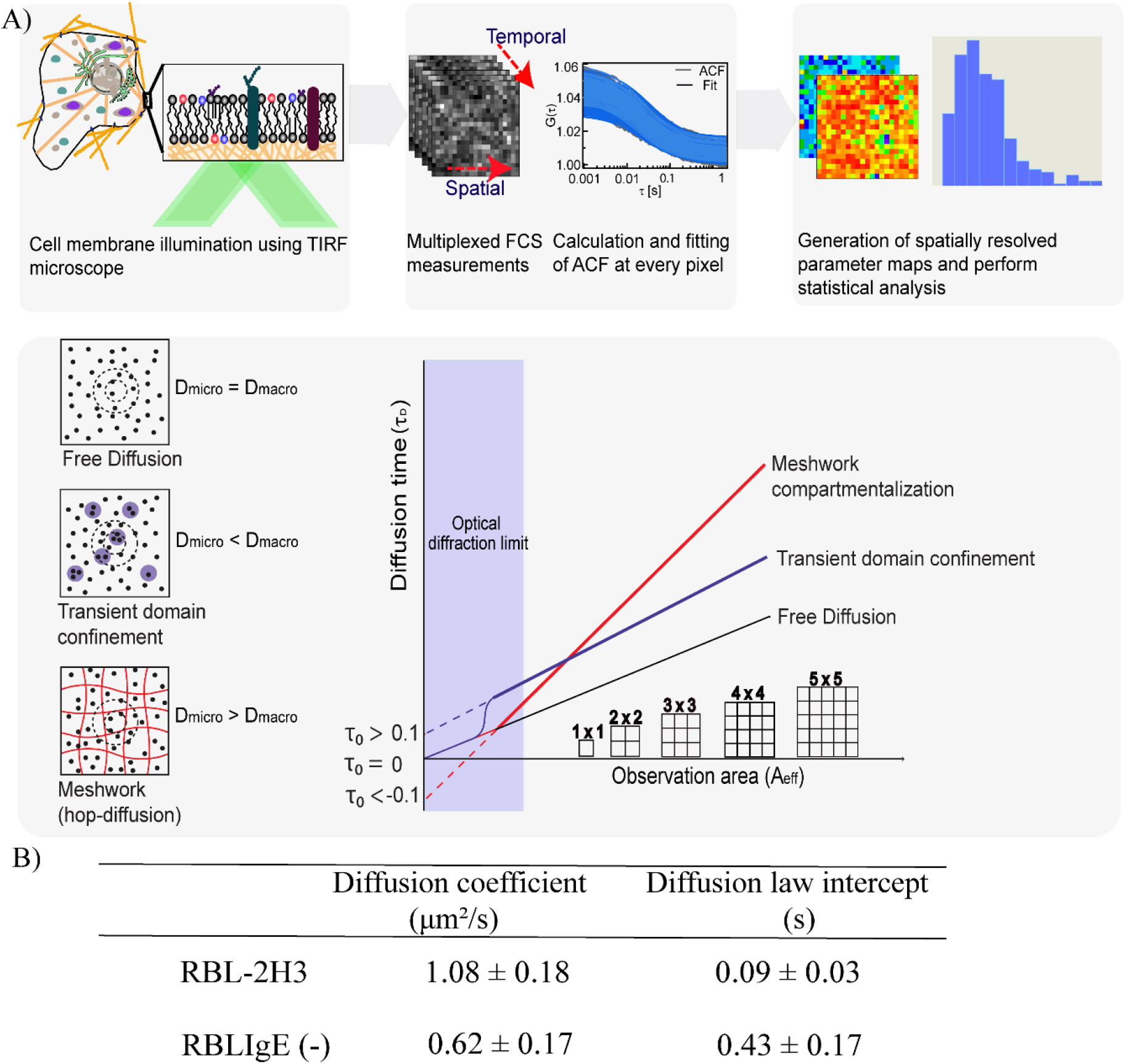
(A) Illustration of a typical imaging total internal reflection fluorescence correlation spectroscopy experiment workflow and imaging FCS diffusion law analysis. (B) Comparison of diffusion coefficients (*D*) and imaging diffusion law plot intercepts (*τ*_*0*_) in RBL-2H3 cells and RBL-IgE(−) cells. Average ± standard deviation (SD) of at least n=3. Refer to Fig *S1* for representative raw data.

### FcεRI knockdown increases DiI-C_18_ confinement in RBL-2H3 cells

DiI-C_18_ exhibits a positive *τ*_*0*_ in RBL-2H3 cells at 25 °C that converges to 0 either if cholesterol is depleted or if the temperature is raised to 37 °C, indicating transient domain trapping. In addition, it showed a weak cytoskeleton dependence in this cell line, indicated by a rise in *τ*_*0*_ when the cytoskeleton was disrupted. In contrast, in CHO-K1 cells, the *τ*_*0*_ is always close to 0 independent of cholesterol and the cytoskeleton^27^. To determine whether the high level of FcεRI is the cause for this diffusive behaviour of DiI-C_18_, we reduced FcεRI levels by siRNA mediated knockdown in RBL-2H3 cells (referred to as RBL-IgE(−)). FcεRIα knockdown was confirmed by western blotting (Fig S2*A*) and staining of cells with a fluorescently labelled FcεRI receptor antibody (Fig S2*B*). Measurements were performed at 37 °C before and after FcεRI knockdown. However, after FcεRI knockdown, the *D* of DiI-C_18_ decreased by 43% (Fig 1*B*) and *τ*_*0*_ increased from a value close to 0 for free diffusion to 0.43 s (Fig 1*B*), implying that DiI-C_18_ is transiently trapped.

### DiI-C_18_ diffusion exhibits a stronger dependence on cholesterol and the cytoskeleton in RBL-IgE(−) cells

Next we probed the sensitivity of DiI-C_18_ diffusion to cholesterol content and cytoskeleton integrity in RBL-IgE**(−)**cell membranes. DiI-C_18_ diffusion was measured before and after drug treatments and results were compared with the DiI-C_18_ diffusion properties measured on CHO-K1 and RBL-2H3 cell membranes.

DiI-C_18_ diffusion on CHO-K1 cells shows changes that are within the margins of error and are not significant (Figure 2 *A, B, C*). In RBL-2H3 cells, cytoskeletal disruption resulted in an increase in *τ*_*0*_ (0.10 s to 0.24 s) and no significant difference in *D*. Cholesterol depletion led to an increase in *D* by 35% but no significant change in *τ*_*0*_. These results suggest that in CHO-K1 cells DiI-C_18_ diffusion is not linked to cholesterol or the cytoskeleton, while in RBL-2H3 cells DiI-C_18_ diffusion is hindered by both cholesterol and the cytoskeleton. Interestingly, in RBL-IgE(−) cells, cytoskeleton disruption resulted in an increase of *D* by 90% and we observed a drop in *τ*_*0*_ from 0.8 s to 0.3 s. Cholesterol depletion caused a drop in *D* by 54%, and an increase in *τ*_*0*_ from 0.38 s to 1 s. Much to our surprise, in RBL-IgE(−) cells, instead of any decrease in the dependence of DiI-C_18_ on cholesterol and cytoskeleton, there was a stronger dependence as indicated by changes in *D* and *τ*_*0*_ (Figure 2 *A, B, C*) upon methyl β cyclodextrin (mβcd) and Latrunculin A (Lat A) treatments. Moreover, the effect of cholesterol and the cytoskeleton on DiI-C_18_ diffusion is the inverse of the trend obtained on RBL-2H3 cells, where DiI-C_18_ diffusion becomes faster upon cholesterol removal, as shown by an increase in *D* and an increase in *τ*_*0*_ upon cytoskeletal disruption. In summary, FcεRI is not responsible for the higher domain fraction of DiI-C_18_ and linking outer leaflet dynamics with the cytoskeleton in RBL-2H3 cells. However, it significantly influences the membrane dynamics.

**Figure 2:**
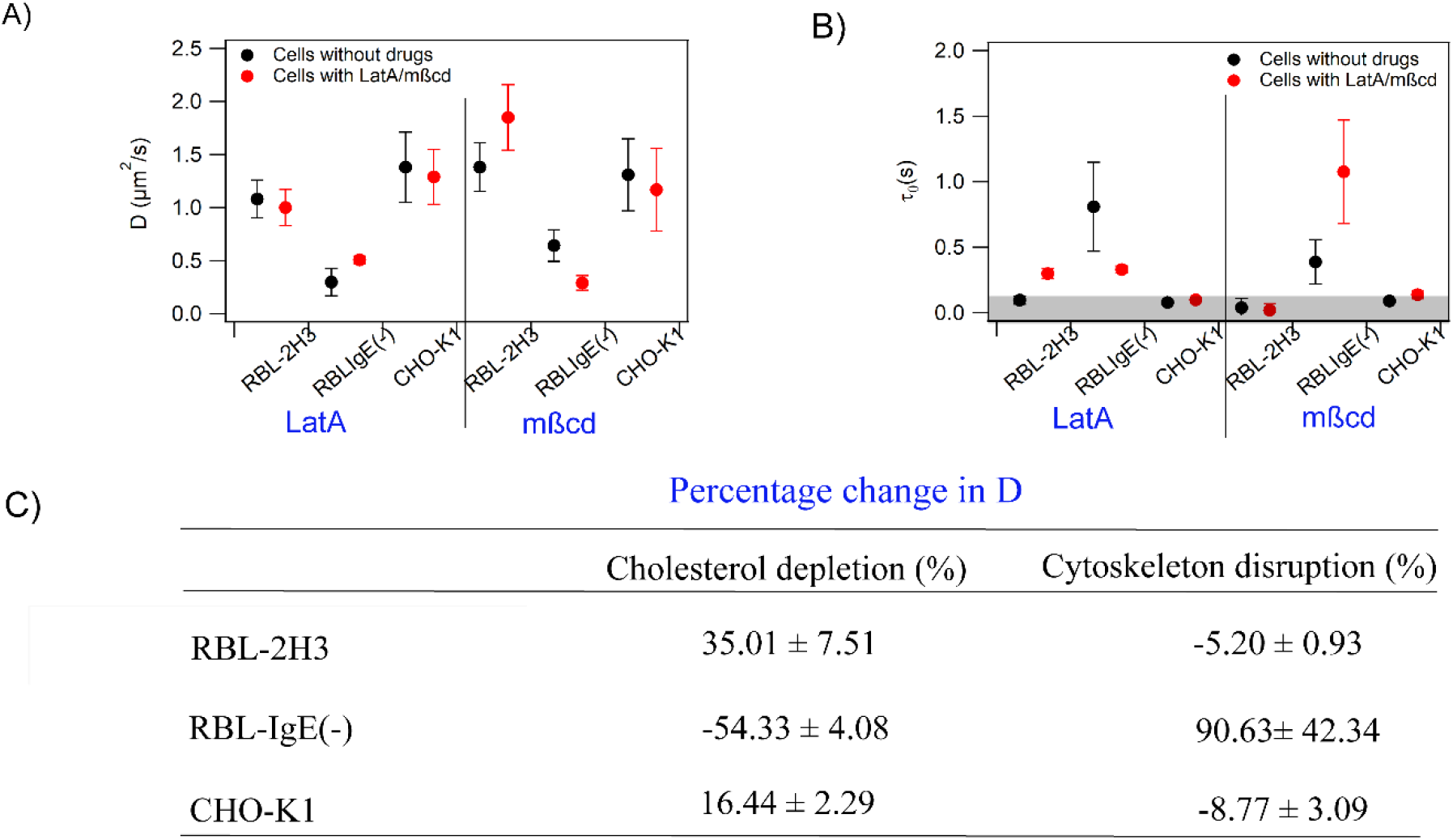
Effect of cholesterol depletion and cytoskeleton disruption in CHO-K1, RBL-2H3 and RBL-IgE(−) cells (A) Comparison of diffusion coefficients (*D*) of DiI-C_18_ in CHO-K1, WT-RBL-2H3, and RBL-IgE(−) cells upon mβcd and Lat A treatments (B) Comparison of imaging FCS diffusion law plot intercepts (τ0) obtained for DiI-C_18_ in CHO-K1, RBL-2H3 cells and RBL-IgE(−) cells upon mβcd and Lat A treatments. Error bars represent standard deviation (SD) of at least n=3. Data represent an average of at least three independent experiments.

### Lipid composition analysis of CHO-K1 and RBL-2H3 plasma membranes using mass spectrometry-based approaches

Next, we analyzed the lipid composition of the plasma membranes of CHO-K1 and RBL-2H3 cells using mass-spectrometry-based lipidomic analysis. The comparison shows markedly different compositions for these two cell types (Table 1). One interesting difference which can potentially explain higher DiI-C_18_ confinement, and E_*Arr*_ comparable to that of domain markers^27^, are the significantly higher levels of sphingolipids, including sphingomyelins and ceramides, in RBL-2H3 cell membranes. As shown in previous studies^31,54–57^, higher sphingolipid content can result in the presence of more domains in RBL-2H3 cell membranes compared to CHO-K1 cells. We therefore followed up on the influence of these lipids on the dynamics of the plasma membrane in the next sections.

**Table 1:** Comparison of plasma membrane lipid composition of in CHO-K1and RBL-2H3 cells by mass spectrometry. Data is represented as average ± standard deviation (SD) of at least 3 replicates. Values are normalized to 100% (concentration/total concentration of detected lipids). Coefficient of variation was less than 20% for about 97% of the analysed lipids in technical quality control samples and less than 30% in the remaining samples (refer Figure *S3*)

| Lipid species | CHO-K1<br>(% lipid/Total PL) | RBL-2H3<br>(% lipid/Total PL) |
| --- | --- | --- |
| PC | 30.84 ± 5.41 | 34.42 ± 11.22 |
| SM | 4.31 ± 2.5 | 13.66 ± 4.56 |
| Cer | 0.17 ± 0.03 | 0.59 ± 0.09 |
| GM3 | 0.30 ± 0.15 | 0.01 ± 0.003 |
| LPC | 0.01 ± 0.004 | 0.10 ± 0.02 |
| LPE | Below detection limit | 0.44 ± 0.03 |
| PI | 20.71 ± 2.12 | 13.11 ± 5.87 |
| PG | 2.18 ± 0.33 | 11.56 ± 3.40 |
| PS | 14.93 ± 1.19 | 6.00 ± 1.56 |
| PE | 26.45 ± 0.53 | 19.63 ± 5.53 |

### Sphingolipid depletion influences the links of DiI-C_18_ diffusion with cholesterol and the cytoskeleton in RBL-2H3 and RBLIgE (−) cells

Since cholesterol and the cytoskeleton influence DiI-C_18_ diffusion in RBL-2H3 and RBL-IgE(−) inversely, they are interesting tools to explore the role of sphingolipids in altering the outer leaflet lipid diffusion properties or DiI-C_18_. First, we depleted sphingolipids by treating cells with myriocin. Myriocin is a common sphingolipid biosynthesis inhibitor^58^. Since myriocin acts on the first step of the pathway, it affects the levels of all sphingolipids. ITIR-FCS experiments were performed to measure DiI-C_18_ diffusion on myriocin treated RBL-2H3 and RBL-IgE(−) cells, followed by cholesterol depletion and cytoskeleton disruption.

In RBL-2H3 cells, myriocin induced sphingolipid depletion altered DiI-C_18_ diffusion as manifested by a 27% lower *D* and a positive *τ*_*0*_ (Figure 3 *A,B*) implying the occurrence of transient domain trapping. In RBL-2H3 cells treated with myriocin, cholesterol depletion causes a 69% drop in *D* and an increase in *τ*_*0*_ from 0.2 s to 1.1 s, indicating an increase of transient domain trapping. This is possibly the case because of non-specific clustering of DiI-C_18_ with FcεRI on RBL-2H3 cell membranes.^44,45^ In myriocin-treated RBL-IgE(−) cells, DiI-C_18_ diffusion shows a 33% rise in *D* and a drop in *τ*_*0*_ from 0.43 s to 0.28 s owing to depletion of sphingolipid domains (Figure 3 *A, B*). Due to a lower FcεRI density in these cells, the effect of FcεRI clustering is less pronounced. In these cells, cholesterol depletion leads to a 44% increase in *D* and a drop in *τ*_*0*_ from 0.31 s to 0.11 s. Therefore, myriocin-treated RBL-IgE(−) cells show cholesterol-dependent confinement of DiI-C_18_ diffusion. The effect of cytoskeletal disruption on DiI-C_18_ diffusion is similar in both RBL-2H3 and RBL-IgE (−) cells. Both cell variants show no significant change in *D* and *τ*_*0*_ (Figure 3 *C,D,E*). This means that reducing the membrane sphingolipid levels decreases the connection between the outer leaflet membrane lipid diffusion and the actin cytoskeleton. These results show that sphingolipids play an important role in orchestrating the cholesterol-cytoskeletal links with DiI-C_18_ diffusion (Figure 3*F*).

**Figure 3:**
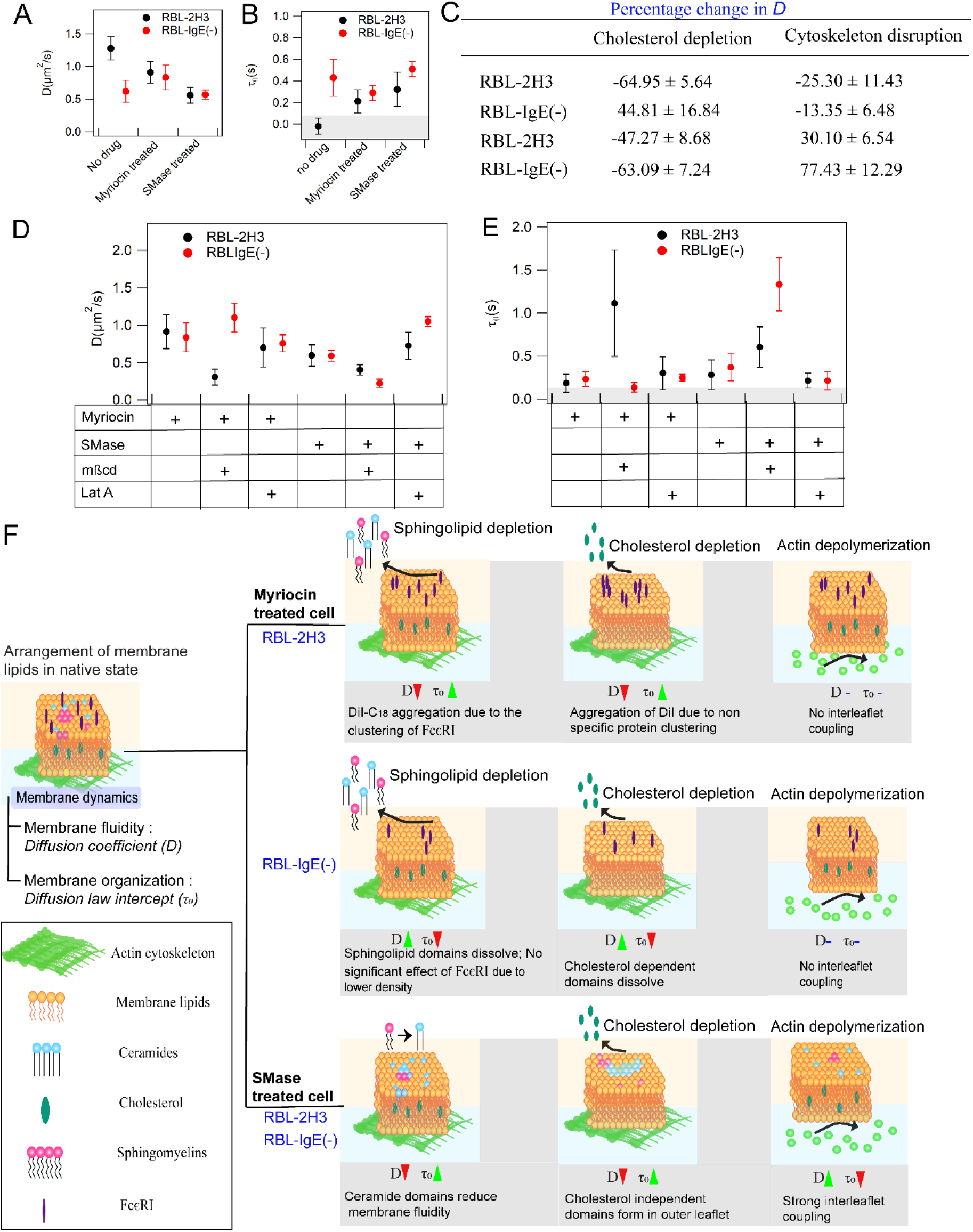
Dependence of outer leaflet lipid diffusion on cholesterol and cytoskeleton upon manipulation of sphingolipid levels in RBL-2H3 and RBL-IgE(−) cells. Comparison of DiI-C_18_ diffusion coefficients (*D*) and (B) FCS diffusion law intercepts (*τ*_*0*_) in wildtype, myriocin and SMase treated cells. Comparison of DiI-C_18_ diffusion in wildtype, myriocin and SMase treated RBL-2H3 cells and RBL-IgE(−) cells upon mβcd and Lat A treatments (C) Percentage change of *D.* Variation of (D) *D* and (E)*τ*_*0*_. Error bars are standard deviation (SD) of at least n=3. Grey area in (B) and (E) represents the region of free diffusion. (F) Schematic illustration of results. FcεRI are not shown for SMase treatment as in this case DiI-C_18_ diffusion is influenced by the change in lipid composition rather than FcεRI density.

### Sphingomyelinase treatment influences the links of DiI-C_18_ diffusion with cholesterol and cytoskeleton in RBL-2H3 and RBLIgE (−) cells

Sphingolipids include ceramides, sphingomyelin and glycosphingolipids. The limited literature on the role of structurally different sphingolipids in regulating biophysical properties of membranes suggests that they have different impact on the membrane dynamics, based on their unique physical properties^59,60^. For instance, ceramides can both order and fluidize the membrane and certain ceramides can form cholesterol independent domains^37,61–63,64^. Additionally, there is evidence that some ceramide species can cause actin cytoskeleton remodelling^65^. This led us to speculate that ceramides could cause the rearrangement of cholesterol-cytoskeletal links. So, to probe the role of ceramides, we treated RBL-2H3 and RBL-IgE(−) cells with sphingomyelinase, which hydrolyzes the membrane sphingomyelin into ceramides^66^, increasing the ceramide content in the plasma membrane. Sphingomyelinase treated cells were subjected to cholesterol depletion and cytoskeleton disruption to analyse the relationship of DiI-C_18_ diffusion with cholesterol and the cytoskeleton under these conditions. Sphingomyelinase treatment in RBL-2H3 cells resulted in a 50% reduction of *D* accompanied with an increase of *τ*_*0*_ from 0 (free diffusion) to 0.3 s (Figure 3 *A,B*) indicative of transiently domain trapped diffusion of DiI-C_18_. Cholesterol depletion in sphingomyelinase treated RBL-2H3 and RBL-IgE(−) cells led to an increase in *τ*_*0*_ (RBL-2H3: 0.28 s to 0.60 s and RBL-IgE(−): 0.33 s to 1.33 s) and a significant decrease in *D* (RBL-2H3: 0.60 μm^2^/s to 0.40 μm^2^/s and RBL-IgE(−): 0.59 μm^2^/s to 0.22 μm^2^/s) (Figure 3 *C,D,E*) indicating elevated confinement of DiI-C_18_ or rise of the ordered domain fraction in the membrane. The cytoskeletal disruption caused a reduction in the confinement of DiI-C_18_ or a drop in the fraction of ordered domains in both cell types as indicated by an increase in *D* (RBL-2H3: 0.61 μm^2^/s to 0.78 μm^2^/s and RBL-IgE(−): 0.59 μm^2^/s to 1.05 μm^2^/s) and a decline in τ0 (RBL-2H3: 0.36 s to 0.15 s and RBL-IgE(−): 0.43 s to 0.20 s) (Figure 3 *C,D,E*). These results show that an increase in ceramides alters the membrane dynamics by reorganizing the lipid domains and the actin cytoskeleton (Figure 3*F*).

### Ceramides influence coupling of outer leaflet lipid dynamics with cholesterol and the cytoskeleton

#### (a) Exogenous ceramide treatment links outer leaflet lipid dynamics with cholesterol

To test if ceramides are sufficient to induce the reorganization of the membrane components, we exogenously treated CHO-K1 cells, where DiI-C_18_ does not show confined diffusion, with those ceramide species which were significantly different in CHO-K1 and the RBL-2H3 cells as determined by lipidomic analysis (Figure 4). These ceramide species are asymmetric long and very long acyl chain ceramides, namely Cer d18:1/16:0, d18:1/18:0, d18:1/24:0, d18:1/24:1. Following the ceramide treatment, cholesterol depletion experiments were performed on the same cells.

**Figure 4:**
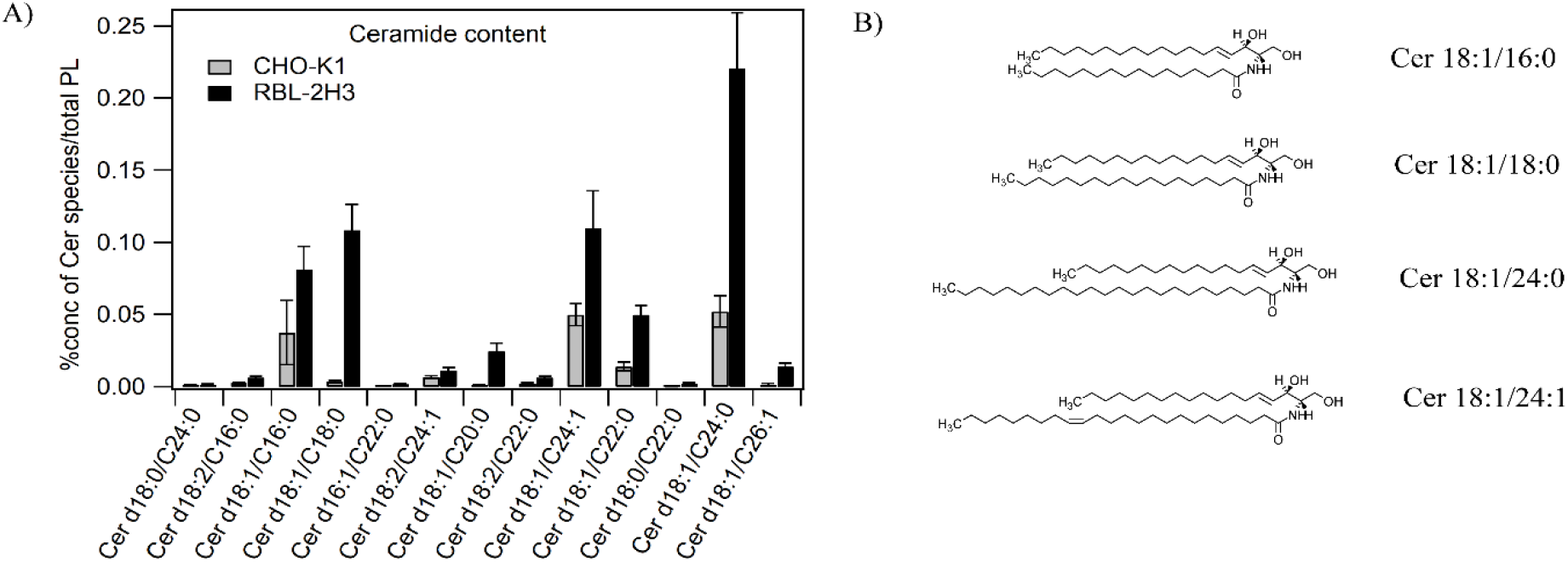
(A) Comparison of plasma membrane ceramide composition in CHO-K1and RBL-2H3 cells analyzed using Agilent 1290 Infinity LC coupled to an Agilent 6495 triple quadrupole mass spectrometer. Error bars are the standard deviation (SD) of n=3. Data were analyzed by Mass Hunter Quantitative Analysis software. Bars represent values normalized to 100% (conc/total phospholipid concentration). (B) Chemical structures of Ceramides (d18:1/16:0, d18:1/18:0, d18:1/24:0, d18:1/24:1).

In untreated CHO-K1 cells, DiI-C_18_ diffusion was not sensitive to cholesterol depletion as shown in Figure 5 *A, B, E*. Treatment of all tested ceramides reduced the *D* of DiI-C_18_ and increased the *τ*_*0*_, and therefore induced confined diffusion of DiI-C_18_. Interestingly, upon cholesterol depletion, Cer d18:1/16:0 and d18:1/24:0 treated CHO-K1 cells showed an increase in the DiI-C_18_ confinement, as indicated by an increase in *τ*_*0*_ and a drop in *D* (Figure 5 *A, B, E*). In the case of Cer d18:1/24:1, cholesterol removal decreased the confinement of DiI-C_18_ in CHO-K1 cell membranes. Upon treatment with Cer d18:1/18:0, cholesterol removal slightly reduced the DiI-C_18_ confinement. Cer d18:1/24:0 treated cells showed maximum sensitivity to cholesterol removal as indicated by a 58% reduction in *D* (0.96 μm^2^/s to 0.34 μm^2^/s) after the treatment. These effects are similar to the results in the RBL-2H3 cells, which show cholesterol hindered DiI-C_18_ diffusion even without the addition of ceramides. When RBL-2H3 cells were additionally treated with Cer d18:1/16:0 and d18:1/24:0, cholesterol depletion increased the DiI-C_18_ confinement, shown by a further increase in *τ*_*0*_ (Figure 5 *C, D, E*). In the case of Cer d18:1/24:1 and d18:1/18:0 treated RBL-2H3 cells, cholesterol depletion lowers the confinement of DiI-C_18_, as shown by a drop in *τ*_*0*_ (Figure 5 *C, D, E*). In RBL-2H3 cells Cer d18:1/24:0 influences membrane dynamics the most (D: 0.76 μm^2^/s to 0.42 μm^2^/s; *τ*_*0*_: 0.30 s to 0.80 s).

**Figure 5:**
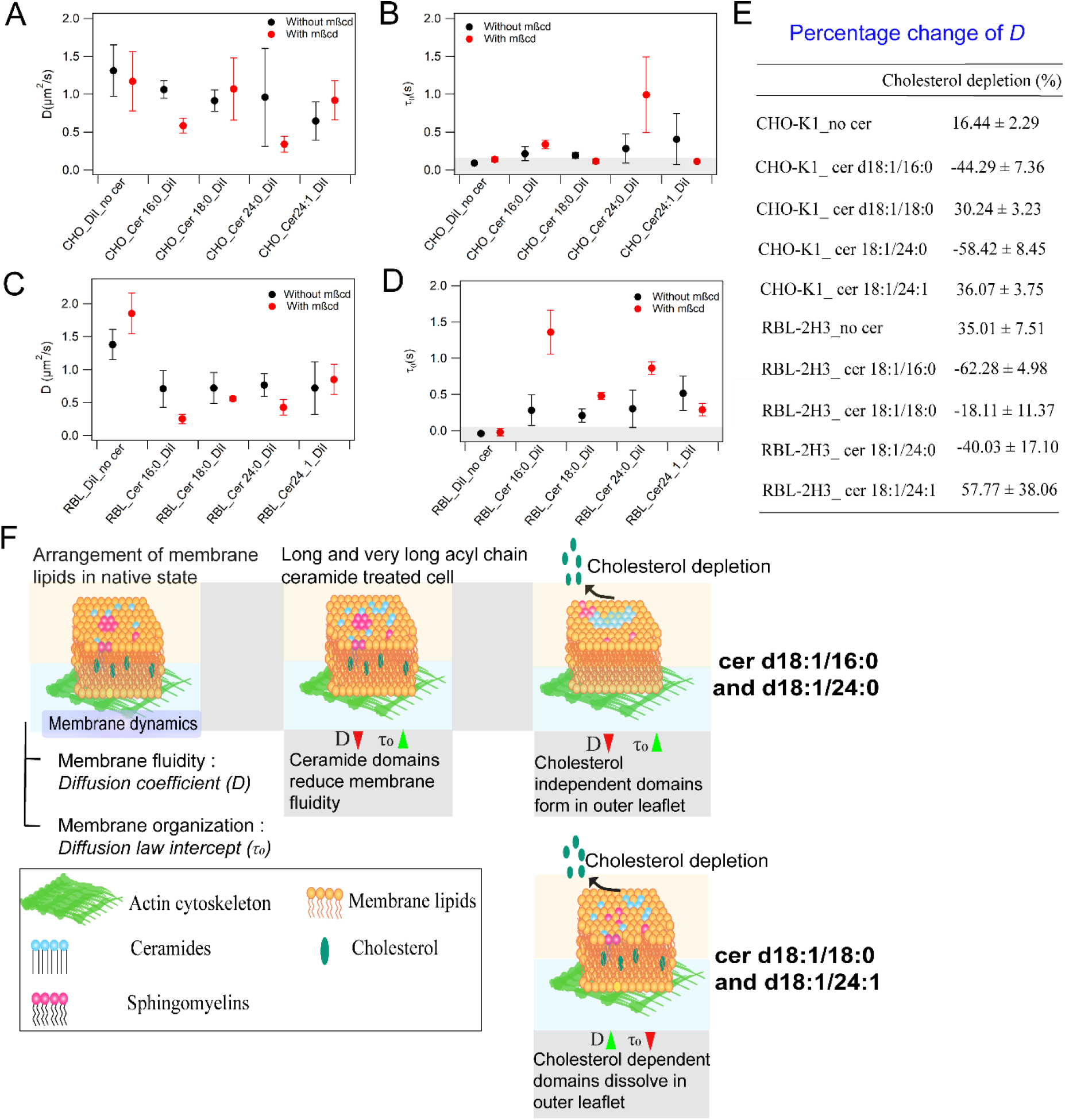
Effect of cholesterol depletion in ceramide (cer d18:1/16:0, cer d18:1/18:0, cer d18:1/24:0 and cer d18:1/24:1) treated CHO-K1and RBL-2H3 cells (A) Comparison of diffusion coefficients (*D*) of DiI-C_18_ in CHO-K1 cells treated with ceramides (B) Comparison of imaging FCS diffusion law plot intercepts (*τ*_*0*_) obtained for DiI-C_18_ in CHO-K1 cells treated with ceramides. (C) Comparison of diffusion coefficients (*D*) of DiI-C_18_ in RBL-2H3 cells treated with ceramides. (D) Comparison of imaging FCS diffusion law plot intercepts (*τ*_*0*_) obtained for DiI-C_18_ in RBL-2H3 cells treated with ceramides. Grey area in (B) and (D) represents the region of free diffusion. E) Percentage change in the diffusion coefficient (*D*) before and after the cholesterol depletion. Error bars are the standard deviation (SD) of at least n=3 (F) Schematic illustration of results.

Our results suggest that, at lower cholesterol concentrations, Ceramides (Cer d18:1/16:0 and Cer d18:1/24:0) can form cholesterol independent domains in the outer leaflet of both CHO-K1 and RBL-2H3 cell membranes (Figure 5*F*).

#### (b) Exogenous ceramide treatment links outer leaflet lipid dynamics with the cytoskeleton

Next, we probed how outer leaflet lipid diffusion connects with the actin cytoskeleton network in cells treated with these ceramides. Following ceramide treatment, cells labelled with DiI-C_18_ were measured before and after cytoskeleton disruption by Lat A. Cytoskeletal disruption reduced the confinement of DiI-C_18_ in CHO-K1 cells treated with Cer d18:1/16:0 and d18:1/24:0 as reflected by an increase in *D* and decreased *τ*_*0*_ (Figure 6 *A, B, E*). Cytoskeletal disruption slightly increased the confinement of DiI-C_18_ in Cer d18:1/24:1 treated cells. Cer d18:1/18:0 showed no significant change in the DiI-C_18_ diffusion on the CHO-K1 cell membrane (Figure 6 *A, B, E*). As observed for cholesterol sensitivity, the effect of cytoskeletal disruption is maximal in the case of Cer d18:1/24:0 treated CHO-K1 cells with a 167% change of *D* (D: 0.76 μm^2^/s to 2.14 μm^2^/s; *τ*_*0*_: 0.27 s to 0.14 s). Cer d18:1/18:0 shows the least effect on the DiI-C_18_ diffusion properties as indicated an insignificant (9%) change in *D*. Similarly, RBL-2H3 cells treated with Cer d18:1/16:0 and d18:1/24:0 show a reduction in DiI confinement upon cytoskeleton disruption, with Cer d18:1/24:0 showing a maximal effect as shown by 154% change in *D* (*D*: 0.27 μm^2^/s to 0.67 μm^2^/s; *τ*_*0*_: 0.88 s to 0.35 s). In Cer d18:1/18:0 and d18:1/24:1 treated cells, DiI-C_18_ does not show any considerable sensitivity to cytoskeleton disruption indicated by less than 20% change in *D* except that in Cer d18:1/18:0 treated RBL-2H3 cells the cytoskeleton disruption causes a slight increase in *τ*_*0*_ (0.45 s to 0.63 (Figure 6 *C, D, E*). In summary, DiI-C_18_ diffusion in CHO-K1 cell membranes becomes sensitive to the cytoskeleton upon ceramide treatment. However, the effect of ceramide treatment on the arrangement of lipid domains and cytoskeleton varies with the ceramide species. These results demonstrate that the addition of long and very long chain acyl chain ceramides into the membrane increases the confinement within the membranes in general and it establishes links between cytoskeleton and the outer leaflet lipid dynamics.

**Figure 6:**
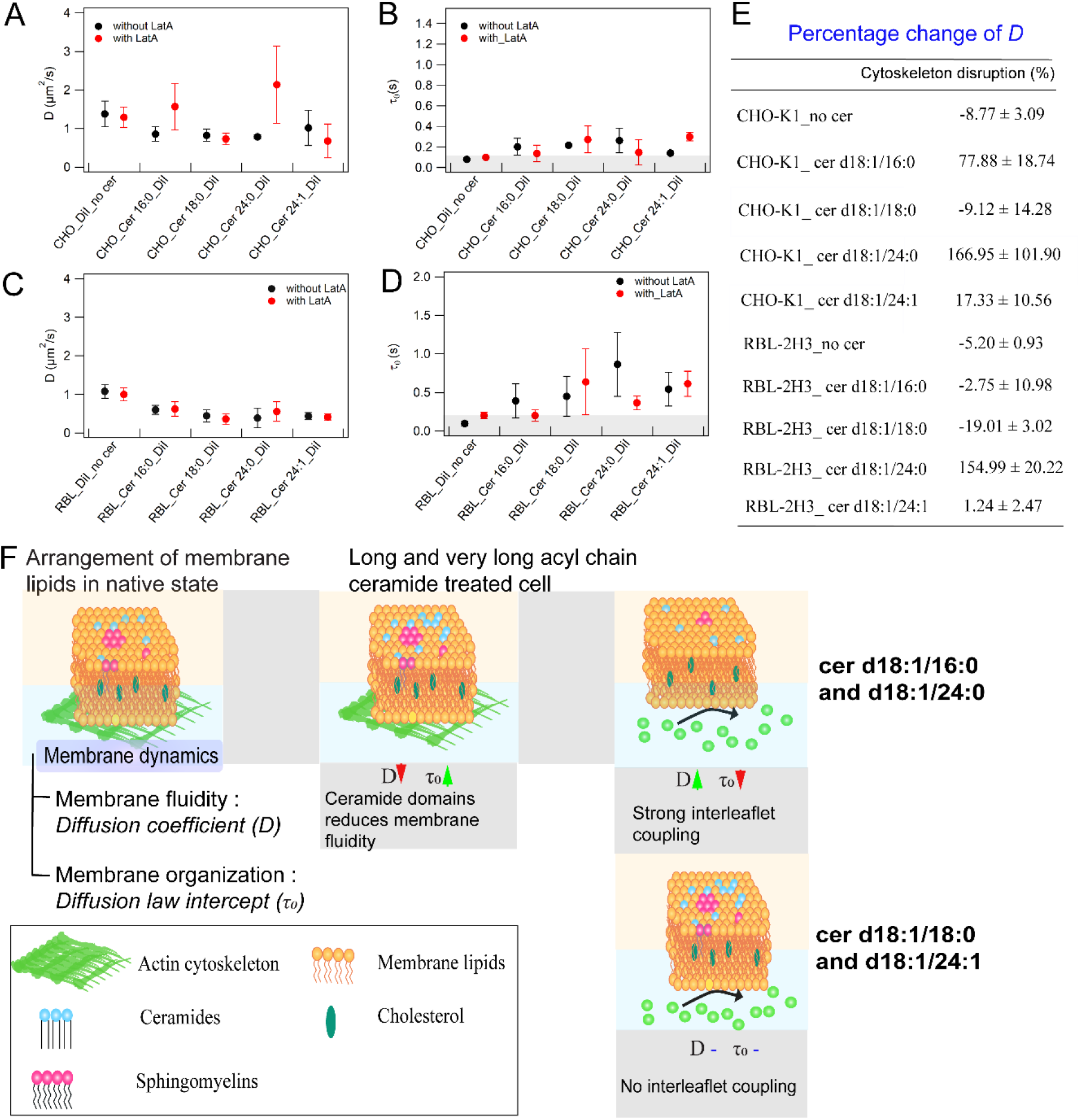
Effect of cytoskeleton disruption in ceramide (cer 18:1/16:0, cer 18:1/18:0, cer 18:1/24:0 and cer 18:1/24:1) treated CHO-K1and RBL-2H3 cells (A) Comparison of diffusion coefficients (*D*) of DiI-C_18_ in CHO-K1 cells treated with ceramides (B) Comparison of imaging FCS diffusion law plot intercepts (*τ*_*0*_) obtained for DiI-C_18_ in CHO-K1 cells treated with ceramides. (C) Comparison of diffusion coefficients (*D*) of DiI-C_18_ in RBL-2H3 cells treated with ceramides. (D) Comparison of imaging FCS diffusion law plot intercepts (*τ*_*0*_) obtained for DiI-C_18_ in RBL-2H3 cells treated with ceramides. Grey area in (B) and (D) represents the region of free diffusion. Error bars are the standard deviation (SD) of at least n=3. (E) Percentage change of *D* for ceramide treated cells before and after the cytoskeleton disruption. (F) Schematic illustration of results.

We next investigated if ceramides can cause any actin cytoskeleton restructuring which could explain the influence of cytoskeleton disruption on DiI-C_18_ diffusion. For this purpose, we treated the CHO-K1 and RBL-2H3 cells expressing lifeact-GFP with the same set of ceramides that have been tested in the previous experiments and performed TIRF microscopy. Lifeact is a 17-amino-acid peptide, which is used in eukaryotic cells to stain filamentous actin (F-actin) structures^67^. It does not interfere with the dynamical changes in actin both *in vitro* and *in vivo*, therefore it allows the visualization of actin dynamics in cells. TIRF microscopy focuses specifically on the cytoskeleton network near the membrane.

The cytoskeletal arrangements in CHO-K1 cells and RBL-2H3 cells are entirely different. At the microscopic level, CHO-K1 (Figure *S4A*) cells showed lifeact-labeled stress fibers while RBL-2H3 (Figure *S5*) cells showed lifeact puncta.

On treating CHO-K1 cells with Cer d18:1/16:0 and Cer d18:1/24:0, the number of filaments decreased, and the length of existing filaments was shorter (Figure *S4 B, C*). On the other hand, there was no quantifiable microscopic difference in the actin cytoskeleton arrangement on treating the cells with Cer d18:1/18:0 and d18:1/24:1. In the case of RBL-2H3 cells, the addition of Cer d18:1/16:0 and d18:1/24:0 decreased the number of lifeact puncta, and the overall intensity was significantly reduced (Figure *S5 A, B*). Cer d18:1/18:0 and d18:1/24:1 treatment in RBL-2H3 cells did not show any significant effect on the actin cytoskeleton rearrangement. These observations suggest that Cer d18:1/16:0 and d18:1/24:0 disorganize the actin cytoskeleton network proximal to the membrane. Cer d18:1/18:0 and d18:1/24:1 do not alter cytoskeleton structure at the microscopic level.

Our results clearly demonstrate that ceramides are sufficient to induce transbilayer coupling in the cell membranes. Cer d18:1/16:0 and d18:1/24:0 restructure the cytoskeleton arrangement and lead to the formation of cytoskeleton induced domains in the outer leaflet of the plasma membrane (Figure 6*F*).

#### (a) Comparison of plasma membrane lipid composition of RBL cells with lower FcεRI density and wildtype RBL-2H3 cells

For biochemical evidence of the membrane ceramide content-dependent sensitivity of outer leaflet lipid dynamics to cholesterol and cytoskeleton, we performed lipid composition analysis of the plasma membrane fractions of RBL-IgE (−) cells and compared it with that of RBL-2H3 cells. In general, no drastic differences were observed in the lipid composition of the two RBL-2H3 cell variants, except the changes observed in the levels of ceramides and lysophospholipids (lysophosphatidylcholines and lysophosphatidylethanolamines) as shown in *Table S1*. The fact that all the major lipids that constitute the plasma membrane remain at similar levels upon FcεRI knockdown indicates that siRNA treatment on RBL-2H3 cells is mild enough to preserve the integrity of the plasma membrane, therefore avoiding artifacts. Upon FcεRI knockdown, there is an increase in LPE levels from 0.44% to 0.67%, and there is a decrease in ceramide content from 0.59% to 0.29%. An inverse correlation of ceramides and lysophospholipids has been previously observed in clinical samples also^68^. These results provide another indication that outer leaflet lipid dynamics is sensitive to the ceramide content of the plasma membrane.

## DISCUSSION

Cell membrane organization and dynamics vary across cell types, and certain cell types tend to show actin induced domain formation on the outer membrane or inter-leaflet coupling ^27,43^. Because of the lack of studies performed on live cells, our understanding of the cell membrane dynamics is still far from complete. Moreover, the role of fine-tuned lipid composition resulting in unique cell type-to-cell type membrane properties is not well understood. In this work, we investigate the factors that can link the outer leaflet lipid diffusion with cytoskeleton and cholesterol in live cell membranes. Our results show that (i) abundant transmembrane proteins interacting with actin may not necessarily link outer leaflet diffusion with the cytoskeleton; (ii) cell membrane ceramide content determine the extent of inter-leaflet coupling and cholesterol dynamics; (iii) specific long and very long acyl chain ceramides induce the formation of actin cytoskeleton-dependent ordered domains and form cholesterol independent domains in the outer leaflet.

### Influence of abundant actin binding transmembrane protein on outer leaflet lipid dynamics in live mammalian cells

Several models have suggested that it is the transmembrane proteins which play a central role in linking outer leaflet organization with the actin cytoskeletal changes^69–71,72^. In a FRAP-based study, it was shown that membrane protein density influences the lateral diffusion of membrane components^16^. Recently, Freeman et al. suggested that transmembrane proteins that interact with both cytoskeleton, via focal adhesion complex proteins, and extracellular matrix exhibit membrane compartmentalization ^71^. Based on these studies, we investigated the role of FcεRI receptors, one of the most abundant transmembrane proteins in RBL-2H3 cells known to interact with the actin cytoskeleton and integrins^50,73,48,49^, in linking outer leaflet lipid diffusion to cholesterol and cytoskeleton. We found that reducing the density of FcεRI receptors on the RBL-2H3 cell membranes results in more confined DiI-C_18_ diffusion and stronger links of outer leaflet lipid diffusion with cholesterol and the cytoskeleton (Fig 1*B*,2). Another interesting observation was that reducing the FcεRI receptor density reverses the effect of cholesterol depletion and actin cytoskeleton disruption on DiI-C_18_ diffusion in RBL-IgE (−) cell membranes, as compared to that in RBL-2H3 cells (Figure 2). This means FcεRI knockdown modulates the membrane organization significantly by increasing the fraction of ordered domains on the outer leaflet of the membrane. Moreover, there is a formation of cholesterol independent domains upon cholesterol removal and a drop in the fraction of ordered domains upon cytoskeleton disruption. This is an interesting observation as in the past only membrane protein diffusion has been shown to be impeded by the actin tethering and not outer leaflet lipid diffusion^74^. These experiments suggest that FcεRI does not limit the molecular mobility in the outer leaflet of the RBL-2H3 plasma membrane. Nevertheless, RBL-IgE(−) cells show elevated coupling of DiI-C_18_ diffusion with the cytoskeleton and cholesterol which indicates that FcεRI knockdown causes membrane restructuring, which could be due to cytoskeletal remodeling, change of membrane composition and membrane lipid reorganization. Our lipidomics results indeed show that modulating the FcεRI density alters the levels of ceramides and lysophospholipids in the plasma membrane (Table *S1*) and triggers membrane reorganization, which is crucial for determining diffusion barriers in the outer leaflet. Despite, the abundance and the interaction of FcεRI with the actin cytoskeleton and focal adhesion complex proteins^45,49,50^ it does not mediate interleaflet coupling. Our results indicate the involvement of other factors that determine the lateral and transbilayer plasma membrane dynamics. These observations necessitate further investigation to define the molecular identity of membrane components that determine the membrane organization and dynamics.

### Long and very long acyl chain ceramides induce cholesterol independent domain formation in the outer-leaflet of live mammalian cell membranes

There is increasing evidence that sphingolipids can modulate cholesterol dynamics in the mammalian cell membranes. Castro et al. showed that in the presence of cholesterol, ceramide domains are solubilized while cholesterol depletion promotes ceramide domain formation^75^. They also suggested that ceramide domain formation is extremely sensitive to the fluid-gel transition of the rest of the lipids. Some sphingolipids can cluster together on their own and form membrane patches^54^. It has been observed that during Gaucher disease, there is an accumulation of glycosphingolipids that alter the properties of the non-raft fraction of the membrane^35^. Due to glycosphingolipid accumulation, there was an increase in the confinement in the non-raft fraction, and they proposed that there is a formation of percolating domains in the outer leaflet. This is in line with previous literature as it has been shown that ceramides can displace cholesterol from rafts and form much bigger and more stable domains^37,76,77,63^. Consistent with these studies, our results show that, upon cholesterol depletion, there is an increase in DiI-C_18_ confinement, and the propensity of cholesterol independent domain formation is sensitive to membrane ceramide content (Figure 5). We observed cholesterol independent domain formation specifically in the case of Cer d18:1/16:0 and d18:1/24:0 treated CHO-K1 and RBL-2H3 cells. Even if a specificity to the ceramide structure was not reported before, C24 SM has been shown to abolish domain formation and induce partitioning of cholesterol in the inner leaflet of GUVs ^78^.

This intricate relationship between cholesterol and ceramides is attributed to the structural similarity of the two lipid types with a small headgroup and high overall hydrophobicity. Interestingly, our observations on live mammalian cells measured under physiological conditions show a similar relationship between cholesterol and ceramide domains.

Recently, it has been shown that in the membrane environment ceramides co-segregate with lysophospholipids due to their large headgroup^79^. The interaction with large headgroup lipids can stabilize ceramide domains. However, it is not clear how lysophospholipids influence membrane dynamics. Our results showed that the levels of lysophospholipids were significantly higher in RBL-2H3 cell membranes relative to CHO-K1 cell membranes (Table 1). Comparing the plasma membrane lipid content of RBL-2H3 with RBL-IgE(−) cells, we observed a change in the levels of ceramide and lysophosphospholipids (Table *S1*). This result indicates that co-segregation of ceramides and lysophospholipids may occur in intact cell membranes.

### Long and very long acyl chain ceramides mediate inter-leaflet coupling in mammalian cell membranes

Our results show that ceramide treated cells, specifically for Cer d18:1/16:0 and d18:1/24:0, exhibit inter-leaflet coupling as actin cytoskeleton disruption diminishes ordered domains in the outer leaflet of the membrane in ceramide treated CHO-K1 cells and RBL-2H3 cells (Figure 6). Moreover, there is evidence of microscopic actin reorganization induced by Cer d18:1/16:0 and d18:1/24:0 treatment (Figure *S4*, *S5*).

There is a growing body of literature demonstrating how ceramides modulate the biophysical properties of the membrane. It is known that several ceramide interacting proteins are intracellular, although most of the sphingolipids reside in the outer leaflet of the membrane^80^. A possible mechanism that can facilitate the interaction of intracellular proteins with ceramides residing in the plasma membrane is the rapid flip-flop of ceramides in the membrane. The spontaneous flip-flop of ceramides has been shown in live cells^81^. It has also been observed that flip-flop of ceramide molecules induces the flip-flop of other membrane lipids^82^.

Recently, it was shown that the generation of ceramides reduces the lateral diffusion of integrins due to increased coupling with the remodeled actin cytoskeletal network^83^. There are several studies which indicated that ceramides induce cortical actin remodelling, that influences biomechanical properties^84^, increase the complex assembly that causes actin polymerization^85^, decrease cell spreading capacity^86^ and influence the interaction between the actin cytoskeleton and ERM proteins^87,65^. Based on our results and the accumulating evidence, we speculate that levels of ceramides can govern transbilayer coupling in cell membranes due to their association with integrins.

Another possible mechanism that can facilitate lipid-mediated transbilayer coupling is interdigitation^21,88^. Mayor and colleagues showed that the change in the lipid content in the outer leaflet could be transduced to the inner leaflet and the transbilayer coupling is important for GPI-APs (located in outer leaflet) nanoclustering^42^ because of the interdigitation that occurs between long acyl chain lipids residing in the two leaflets. Our observations clearly indicated that the presence of long and very long acyl chain ceramides induces the transbilayer coupling in the membrane. We observed the maximum change in outer leaflet lipid diffusion in cells treated with Cer d18:1/24:0 (Figure 5,6).

Conceivably, the interaction of ceramides with the proteins associated with the focal adhesion complex, their ability to rapidly flip-flop and the interdigitation tendency of long acyl chains together are responsible for establishing the connection of the outer leaflet with the actin network organization, which in turn controls the transbilayer membrane properties and diffusion properties of lipids in the outer leaflet of the membrane.

### Alterations in membrane dynamics are dependent on ceramide structure

Slight structural differences conferred alteration in the membrane behaviour of ceramide treated cells, some triggering more dramatic changes in the membrane biophysical properties, as shown by our results (Figure 5,6) and previously published work^65,89,5^. There is no direct correlation between the acyl chain length and induced transbilayer-coupling. In addition to the chain length, the degree of saturation also plays a crucial role in determining the effect of a lipid in the membrane dynamics, as demonstrated by the difference in the action of Cer d18:1/24:0 and Cer d18:1/24:1 on DiI-C_18_ diffusion.

The ceramide species tested in this study are long and very long acyl chain ceramides, and they all reduced the diffusion coefficient and increased the confinement in the outer leaflet of the membrane. Our results also showed that different ceramide species have very specific microscopic effects on the cytoskeleton organization in the membrane which correlates well with the particular trends that we have observed for cholesterol depletion and cytoskeletal disruption on DiI-C_18_ diffusion (Figure S4, S5). Notably, ceramide species that cause microscopic disorganization of the cytoskeleton network are the ones that induce domain formation in response to cholesterol depletion and decrease the membrane order upon cytoskeleton disruption. This might be a crucial step in the remodeling of membrane dynamics by ceramides.

The mammalian cell membrane lipidome has a considerable amount of C16 and C24 sphingolipids. Perturbation of membrane ceramides is associated with specific pathologies such as inflammation, type 2 diabetes, obesity, cancer to name a few. Given our results by lipidomics analysis and the determination of membrane dynamics and organization by ITIR-FCS, there is a possibility that ceramide-induced changes in the membrane dynamics can be correlated with certain pathophysiological conditions.

## CONCLUSION

Diffusion in cell membranes is influenced not only by the lipid and protein composition but also by the asymmetries of the outer and inner leaflets of the membrane as well as the cytoskeleton that can couple to various cell membrane components. Here, we measured the diffusion of DiI, an outer lipid membrane marker in RBL-2H3 and CHO-K1 cells to investigate how proteins and lipids influence the coupling between cytoskeleton and the outer leaflet of the plasma membrane. These cells were chosen as DiI-C_18_ shows free diffusion in CHO-K1 but confined diffusion in RBL-2H3 cells. We demonstrate that for RBL-2H3 cells the abundant FcεRI transmembrane receptors are not the cause of DiI-C_18_ confinement but rather are a factor to limit its confinement. An analysis of lipid composition between RBL-2H3 and CHO-K1 cells indicated specific differences in ceramides (C16 and C24) between the cells, which turned out to be the defining factor in DiI-C_18_ confinement and its coupling to the cytoskeleton and cholesterol. This was shown by exogenous treatment of CHO-K1 with long and very long acyl chain ceramides, which resulted in remodeling of the membrane and in diffusion characteristics similar to those observed in RBL-2H3 cells. Notably, ceramides d18:1/16:0 and d18:1/24:0 are sufficient to promote the formation of cholesterol-independent ceramide domains and interleaflet coupling in the plasma membrane. This study establishes that membrane ceramide content can remarkably remodel the membrane organization and can be a key factor determining the transbilayer connections in the membrane.

## MATERIAL AND METHODS

### Lipids

Ceramide d18:1/16:0, ceramide d18:1/18:0, ceramide d18:1/24:0 and ceramide d18:1/24:1 lipids were purchased from Avanti Polar Lipids (Alabaster, AL). Ceramides were solubilized in ethanol. DiI-C_18_ (Molecular Probes, Invitrogen, Singapore) stock solution was prepared in Dimethyl sulfoxide (DMSO; Sigma-Aldrich, Singapore).

### Cell culture, transfection, and staining

Cell lines – Chinese hamster ovary (CHO-K1) and rat basophilic leukemia (RBL-2H3) – were purchased from ATCC (Manassas, VA). Cells were cultured in DMEM medium (Dulbecco’s Modified Eagle Medium; Invitrogen, Singapore), with 10% FBS (fetal bovine serum; Invitrogen, Singapore) and 1% PS (penicillin and streptomycin; PAA Austria) at physiological temperature (37 °C) in a 5% (v/v) CO_2_ humidified incubator chamber. To ensure that the cells were in their native state, experiments have been performed on cells with passage number lower than 20. For measurements, cells were seeded in mattek dishes with 35 mm (MatTek Corporation, Ashland, Massachusetts, United States). Prior the cell-culture reagents were pre-heated in water bath adjusted to 37 °C.

The stock DiI-C_18_ solution (Molecular Probes, Invitrogen, Singapore) was prepared in DMSO. For DiI staining, stock solution was vigorously vortexed and was diluted in HBSS (Hank’s Balanced Salt Solution; Invitrogen, Singapore) to a final concentration of 100 nM. The cells were incubated with the working solution of DiI-C_18_ at 37 °C for 15-20 minutes. After incubation, the cells were washed with imaging medium (DMEM with no phenol red (Invitrogen, Singapore) and 10 % FBS) at least thrice and then the cells were used for imaging.

Lifeact-GFP plasmid was a kind gift from Dr. Min Wu (CBIS, NUS). Plasmids transfections into the cells was done using the Neon^®^ Transfection System (Invitrogen, Singapore) according to the manufacturer’s manual containing cell culture medium (DMEM + 10% FBS). Post 20 to 48 hours of incubation, the cells were washed with imaging medium twice and imaged in the fresh imaging medium.

### Drug treatments

Methyl beta-cyclodextrin (mβCD) and Latrunculin A (Lat A) were obtained from Sigma-Aldrich (Singapore), and were solubilized in phosphate-buffered saline (PBS; Fluka Biochemicals, Singapore). They were diluted further with the imaging medium to prepare final concentrations of 3 mM and 3 μM, respectively. Myriocin (Myr) and sphingomyelinase were purchased from Sigma-Aldrich (Singapore) and were dissolved in phosphate-buffered saline (PBS; Sigma-Aldrich, Singapore). Further, they were diluted with the imaging medium to make the required final concentrations. The final concentration of myriocin used was 2 μM. The final working concentration of sphingomyelinase was 0.0025 U/mL.

### FcεRI knockdown in RBL-2H3 cells using FcεRIα siRNA treatment

For FcεRIα knockdown in RBL-2H3 cells, cells were treated with sequence-specific predesigned siRNA (Ambion, Singapore). 2 nM siRNA was electroporated in approximately 10^6^ cells, and then the cells were plated on 35 mm uncoated dishes (MatTek Corporation, US). The cells were incubated at 37 °C for 48 hours before imaging. Western blotting verified the effect of siRNA treatment. The primary antibody used was anti-rabbit FcεRIα (R-180): sc-98245 (Santa Cruz Biotechnology, Inc, Germany) and secondary antibody was anti-rabbit IgG-HRP: sc-2357. β-actin was used as the loading control. siRNA mediated knockdown was further confirmed by imaging cells labelled with Alexa Fluor 488 anti-human FcƐRIɑ antibody AER-37 (CRA-1) (BioLegend Products, Singapore).

### Total internal reflection imaging and imaging fluorescence correlation spectroscopy set up, data acquisition and data analysis

#### Imaging Total internal reflection fluorescence correlation spectroscopy (ITIR-FCS)

FCS measurements were performed on an objective type TIRF microscope (IX-71, Olympus, Singapore) with a high NA oil immersion objective (PlanApo, 100×, NA 1.45, Olympus, Singapore). A 532 nm laser (Cobolt Samba, Sweden) coupled into the microscope by a combination of two tilting mirrors was used as the excitation source. The light reflected by a dichroic mirror (Z488/532RPC, Semrock) is focused on the back focal plane of the objective. Subsequently, the light is total internally reflected at the glass-water interface by adjusting the incident angle of the excitation beam by the same combination of tilting mirror. For our experiments, the laser power used was 0.6-1mW. The fluorescence originating from the samples was reflected through the same objective followed by transmission through the same dichroic mirror. Then the fluorescence was filtered by an emission filter (Z488/532M, Semrock). Lastly, the fluorescence was imaged on the CCD chip of a cooled (−80 °C), back-illuminated EMCCD camera (Andor iXON 860, 128×128 pixels, Andor technology, US). The software used for data acquisition is Andor Solis for imaging (version 4.18.30004.0 and 4.24.30004.0). The pixel side length of the CCD chip in the device is 24 μm corresponding to a pixel side length of 240 nm in the sample plane. The camera was operated in the kinetic mode, and baseline clamp was used to minimize the baseline fluctuations. The readout speed was 10 MHz with 4.7x maximum analog-to-digital gain and 25 μs vertical shift speed. An EM gain of 300 was used for most imaging experiments.

The fluorescence signal was concurrently recorded from a 21×21 pixels region of interest (ROI) in the form of a stack of 50,000 frames with a time resolution of 1 ms. The data was saved as 16-bit Tiff file. The temporal intensity trace from each pixel was autocorrelated using multi-tau correlation scheme using a FIJI plug-in ImFCS 1.49, an home-written software which is provided at this link http://www.dbs.nus.edu.sg/lab/BFL/imfcs_image_j_plugin.html] to generate autocorrelation functions (ACF)^90^. To circumvent artefact due to bleaching, the data corrected using a 4^th^ order polynomial function. The ACF for each pixel was individually fitted with the following one-particle model for diffusion using the same software.

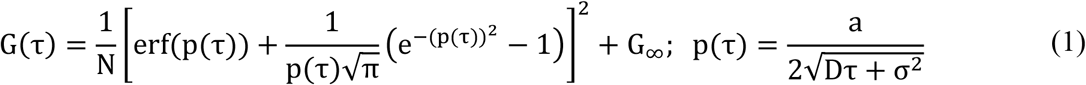

Here *G(τ)* represents the ACF as a function of correlation time (*τ*) and *N*, *a*, *D* and σ are the number of particles per pixel, pixel side length, diffusion coefficient and standard deviation of the Gaussian approximation of the microscope point spread function (PSF) respectively. *G*_∞_ is the convergence value of the ACF at long correlation times.

Fitting of ACFs with theoretical models yields *D* and *N*. Since measurements are performed over a whole region of interest parallelly, diffusion coefficient (*D*), and the number of particles (*N*) maps are spatially resolved^90^. In ITIR-FCS, the data are represented as mean ± standard deviation (SD). The SD is obtained from the measurements over 441 pixels per experiment. The SD of an ITIR-FCS measurement is an indicator of both measurement variability and the lateral heterogeneity of the membrane. All the measurements were performed at 37 °C.

### The FCS Diffusion Laws implemented in ITIR-FCS

For probing the sub-resolution plasma membrane organization, FCS diffusion law analysis is done on data acquired in an ITIR-FCS experiment. With this analysis, one can determine if a particle is exhibiting free diffusion or is hindered by the trapping sites such as membrane domains^51^. This is achieved by plotting the spatial dependence of the diffusion time of the labelled molecules on the observation area. For a freely diffusing particle, the time a particle takes to diffuse through an area is directly proportional to the area (A_eff_). Hence, these plots are fitted to a straight line which is mathematically expressed as:

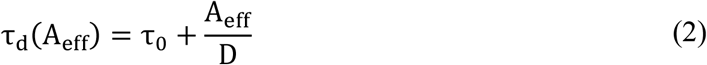

Where *τ*_0_ is the FCS diffusion law intercept. To get sub-resolution information, the diffusion law plot is extrapolated to zero and y-intercept (*τ*_0_) is used as a determinant of membrane organization. For a freely diffusing particle, the diffusion time scales linearly with the observation area and the y-intercept is zero. In case of a domain partitioning, the relationship between the diffusion time of the molecules and the observation area deviates from linearity and y-intercept is positive. To perform FCS Diffusion Law analysis, the same set of data that is acquired in an ITIR-FCS experiment is used. Post-acquisition pixel binning (1×1 to 5×5) followed by convolution with the PSF of the microscope system is performed to obtain variable observation areas (*A*_eff_). The *A*_eff_/*D* is plotted against *A*_eff_, and the plot is fitted to a line with the standard error of mean (SEM) weighted equation (2) to obtain the y-intercept *τ*_0_. The typical margin of error on cell membranes is ±0.1 and thus intercepts in that range are indistinguishable for free diffusion. Only intercepts greater than 0.1 can be attributed to domain trapping in our setup^91^.

### Plasma membrane isolation and lipid extraction

The protocol used for plasma membrane isolation is adapted from Cohen et al. and is slightly modified to increase the yield^92^. For plasma membrane extraction approximately 10^7^ cells of each cell line were taken. Cells were attached on the lysine coated cytodex3 beads (Sigma-Aldrich, Singapore) by incubating 10^7^ cells overnight with 4 mL beads. Cells were maintained at 37 °C with slow shaking. Cells attached on beads were collected via 37 μm reversible strainer, large (Stem cell technologies, Singapore). The media was discarded, and cells attached to beads were collected in 1X PBS. 1X PBS was aspirated (without aspirating beads), and the beads were resuspended in the attachment buffer (220 mM Sucrose, 40 mM sodium acetate, pH 5.0) at room temperature and were incubated for 5 minutes. Then the attachment buffer was aspirated out, and beads were resuspended in hypotonic buffer (10mM Tris-HCl, pH 8.0). After 5 minutes incubation, beads were washed with hypotonic buffer three times, and the beads were sonicated. The sonication strength was optimized so that it doesn't damage the beads. Then the beads were washed at least two times with hypotonic solution. 700 μL of beads were resuspended in 500 μL of cold butanol/methanol (1:1) spiked with 5 μL of SPLASH LIPIDOMIX Mass Spec Standard (330707) and 5 μL of Ceramide/Sphingoid Internal Standard Mixture I (LM6002), from AVANTI. It was incubated at 4 °C for 2 hours with shaking. To separate beads from the solvent, beads were centrifuged at 10,000 rpm for 15 minutes at 4 °C. Then the collected supernatant was evaporated, and it was reconstituted in 100 μL of solvent without the standards. The samples were then subjected to liquid chromatography electrospray ionization tandem mass spectrometry (LC-MS) using Agilent 1290 Infinity LC coupled to an Agilent 6495 triple quadrupole mass spectrometer.

### Liquid chromatography-mass spectrometry of cell extracts: Instrumentation, data acquisition and data analysis

An Agilent ZORBAX RRHD Eclipse plus C18, 95 Å, 2.1 x 100 mm, 1.8 μm UPLC column was used for LC separation at 40 ⁰C. Mobile phases A (60% water and 40% acetonitrile with 10 mM ammonium formate) and B (10% acetonitrile and 90% isopropanol with 10 mM ammonium formate) were used to create the following gradient: 20% B at 0 min to 60% B at 2min; 100% B at 7 min; 100%B at 9 min, 20%B at 9.01 min and 20%B at 10.8 min. The flow rate was set at 0.4 mL/min, and 2 μL of sample were injected. The capillary voltage and nozzle voltage were set at 3,500 V and 500 V, respectively. The drying gas and sheath gas temperatures were maintained at 200 °C and 250 °C, respectively. The drying gas and sheath gas flow rates were 12 L/min and 12 L/min, respectively. Data was exported and analyzed with Mass Hunter Quant Software (Agilent) and the lipids quantified after normalizing for each class-specific internal standard spiked before lipid extraction. A correction for the initial amount of cells was also applied.

For the technical validation of the lipidomics dataset, we analysed internal standards concentration, three biological replicates for each sample and quality control samples (QC). Technical reproducibility of the data is ensured by calculating the coefficient of variation of lipids analysed across QC samples and by maintaining a signal-to-noise ratio of at least 10.

### Image analysis and quantification

To quantify the filament number, length and lifeact intensity within each cell, the Imaris software package “Cells” module (Bitplane, USA Version 8) was used to batch process the cells.

