## Supplementary file 1 for "Long acyl chain ceramides govern cholesterol and cytoskeleton dependence of membrane outer leaflet dynamics"

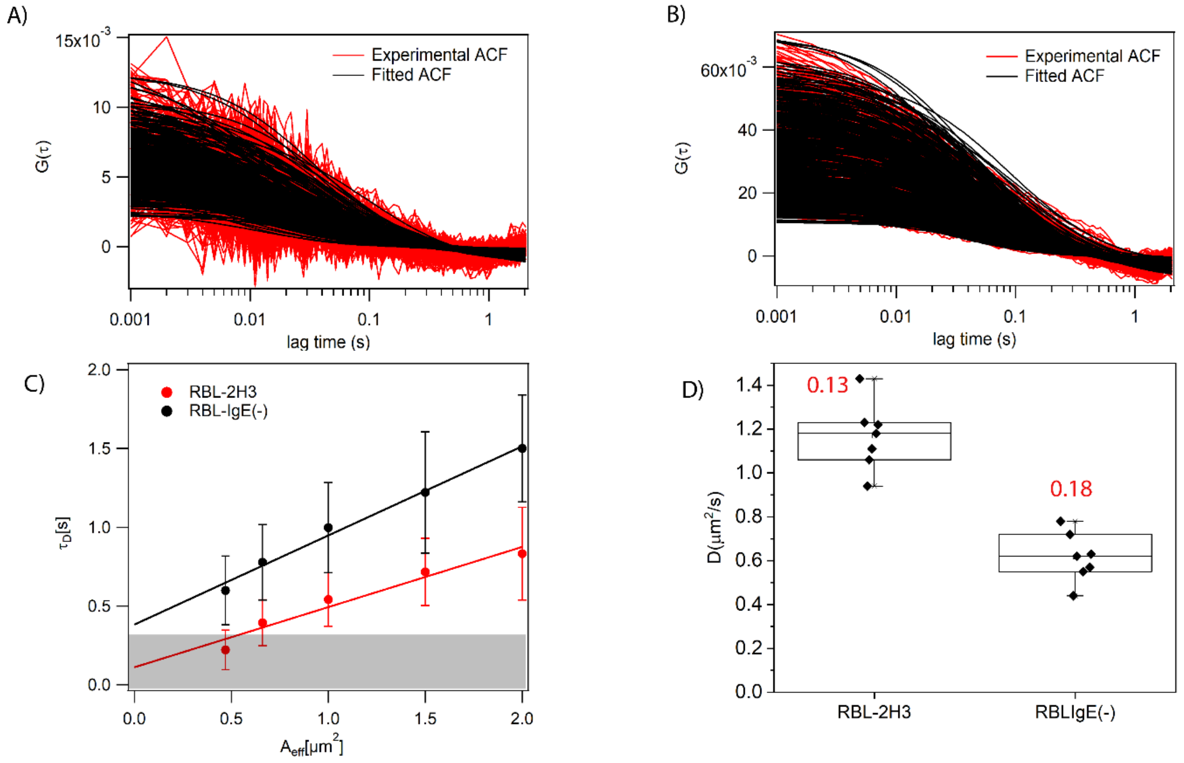


**Figure S1:** (A) Representative autocorrelations for DiI diffusion in RBL-2H3 and (B) RBL-IgE(-) cell membranes (C) Representative FCS diffusion law plot for DiI in RBL-2H3 and RBL-IgE(-) (D) Diffusion coefficient of DiI-$C_{18}$ stained on RBL-2H3 and RBL IgE(-) cell membranes. Numbers on the top of the data are coefficient of variation for the corresponding cell type. Each data point is an average of 441 diffusion coefficients obtained from 441 autocorrelation functions (ACF). Therefore, N = 3087 ACFs for each cell type.

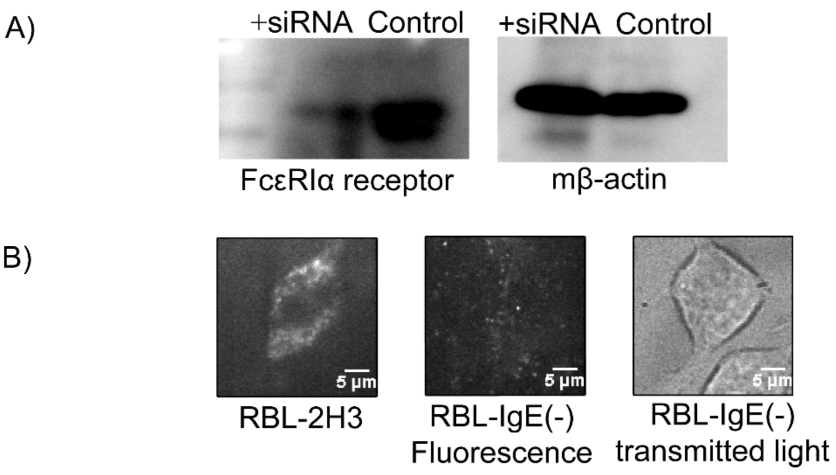


**Figure S2:** Knockdown FcεRI in RBL-2H3 (A) Western blot of FcεRIα siRNA transfected cell compared with RBL-2H3 cells. (B) Images of Alexafluor488 conjugated FcεRI antibody labeled RBL-2H3 cells, and RBL-IgE(-) cells.

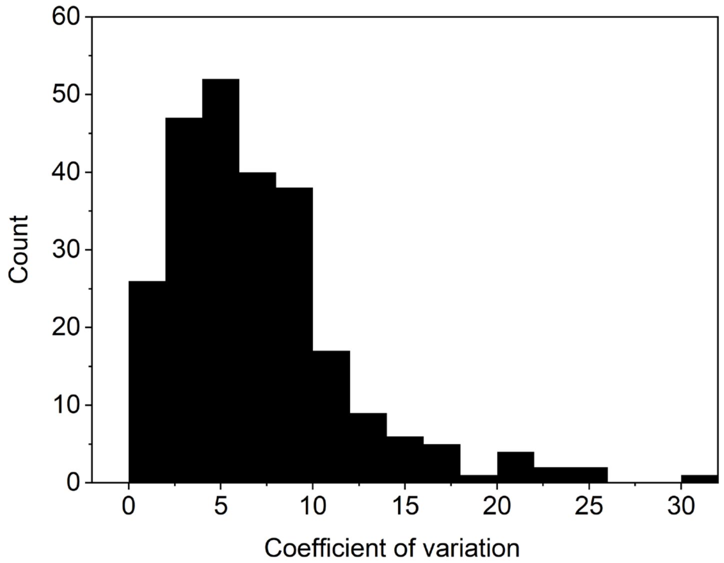


**Figure S3:** Histogram shows the distribution of coefficient of variation (%) for all the lipids analysed in the QC samples while comparing CHO-K1 and RBL-2H3 plasma membranes.

| **Lipid species** | **RBL-2H3 (% lipid/Total PL)** | **RBLIgE(-) (% lipid/Total PL)** |
|---|---|---|
| PC | 34.42 ± 11.22 | 39.41 ± 6.67 |
| SM | 13.66 ± 4.56 | 10.68 ± 1.62 |
| Cer* | 0.59 ± 0.09 | 0.28 ± 0.05 |
| GM3 | 0.01 ± 0.003 | Below detection limit |
| LPC | 0.10 ± 0.02 | 0.14 ± 0.01 |
| LPE* | 0.44 ± 0.03 | 0.67 ± 0.05 |
| PI | 13.11 ± 5.87 | 8.27 ± 0.93 |
| PG | 11.56 ± 3.40 | 20.55 ± 7.12 |
| PS | 6.00 ± 1.56 | 4.77 ± 1.07 |
| PE | 19.63 ± 5.53 | 13.15 ± 1.97 |

**Table S1:** Comparison of plasma membrane lipid composition of in RBL-2H3 and RBLIgE(-) cells analyzed by mass spectrometry. Error bars are the standard deviation (SD) of at least three replicates. Data were analyzed by Mass Hunter Qualitative Analysis software. Values are normalized to 100% (conc/total phospholipid concentration). p-values were calculated for each lipid class, *< 0.05, from 3 independent experiments.

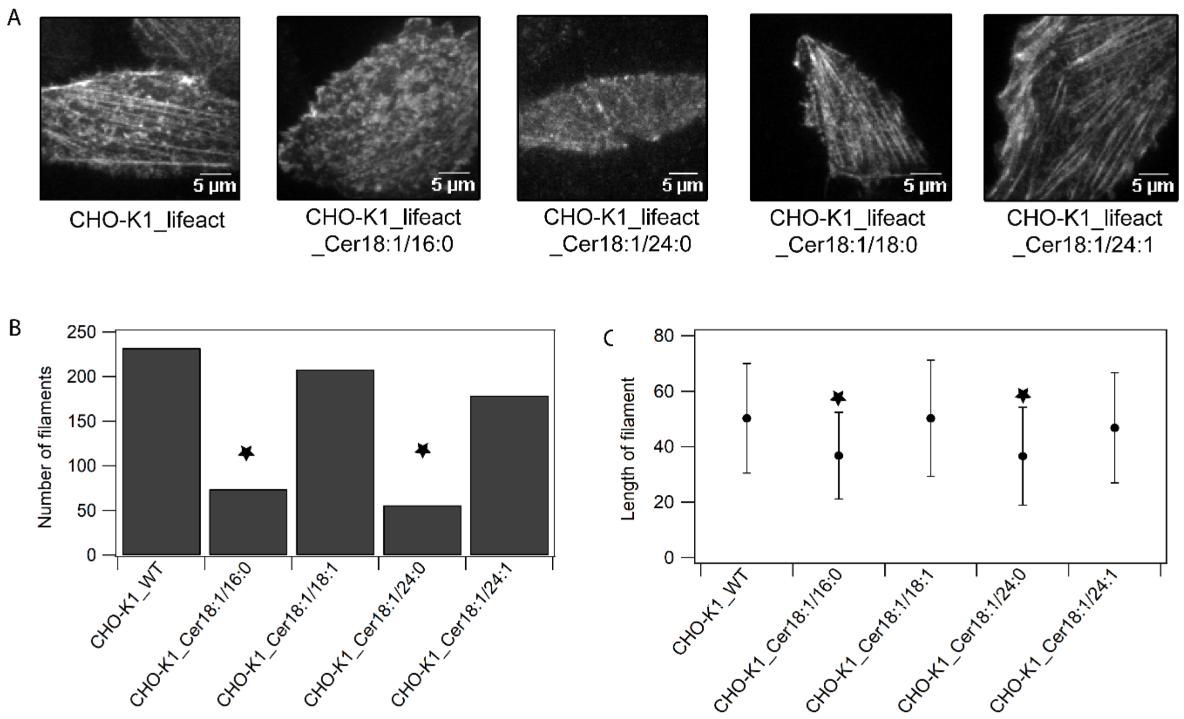


**Figure S4:** Effect of exogenous treatment of ceramides (d18:1/16:0, d18:1/18:0, d18:1/24:0 and d18:1/24:1) on the actin cytoskeleton dynamics in CHO-K1 cells probed by Lifeact-mGFP (A) Representative TIRF images to compare the microscopic actin cytoskeleton organization in CHO-K1 cells upon treatment with various ceramide lipid species. (B) Comparison of the number of filaments in CHO-K1 cells treated with ceramides and the CHO-K1 cells (n=20) (C) Comparison of the length of actin fiber length in CHO-K1 cells treated with ceramides and in CHO-K1 cells. Error bars are the standard deviation (SD). p-values were compared to the respective untreated cells, *< 0.05, n = 20 from 3 independent experiments.

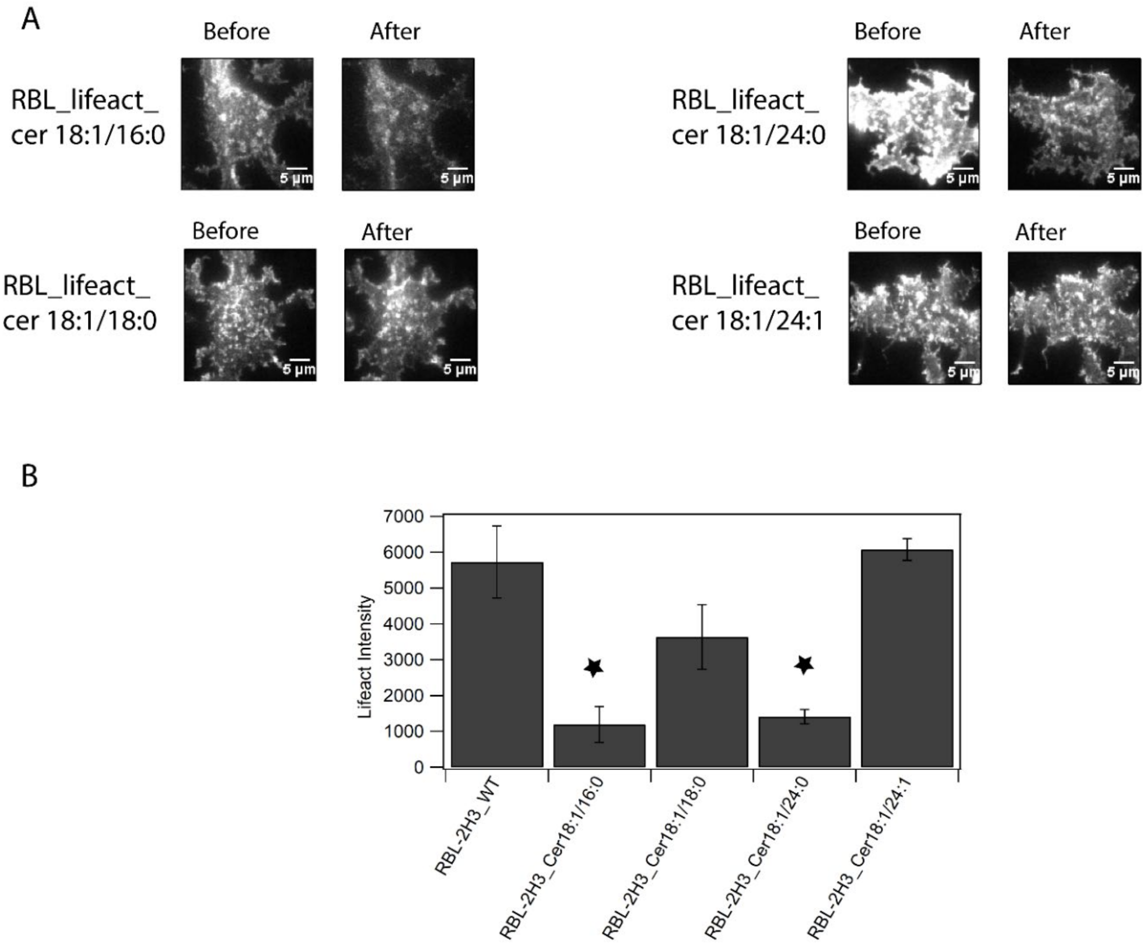


**Figure S5:** Effect of exogenous treatment of ceramides (d18:1/16:0, d18:1/18:0, d18:1/24:0 and d18:1/24:1) on the actin cytoskeleton dynamics in RBL-2H3 cells probed by Lifeact-mGFP (A) Representative TIRF images to compare the microscopic actin cytoskeleton organization in RBL-2H3 cells upon treatment with various ceramide lipid species (before and after for the same cell). (B) Comparison of the intensity of Lifeact-mGFP in RBL-2H3 cells treated with ceramides and the RBL-2H3 cells (n=10). Error bars are the standard deviation (SD). p-values were compared to the respective untreated cells, *< 0.05, n = 10 from 3 independent experiments.
